# A bead-based method for the removal of the amino acid lysine from cell-free transcription-translation systems

**DOI:** 10.1101/2020.01.15.907360

**Authors:** Marc Finkler, Ömer Kurt, Florent Grimm, Philip Hartz, Albrecht Ott

## Abstract

Cell-free transcription-translation systems are a versatile tool to study gene expression, enzymatic reactions and biochemical regulation mechanisms. Because cell-free transcription-translation systems are often derived from cell lysates, many different substances, among them amino acids, are present. However, experiments concerning the incorporation of non-canonical amino acids into proteins require a system with negligible amounts of canonical analogs. Here we propose a two-step method for the removal of residual free lysine in an all *Escherichia coli*-based cell-free expression system. The first step consists of the expression of a high-lysine dummy protein. The second step consists of direct removal via binding between lysine and DNA. The presented method is an efficient, fast and simple way to remove residual lysine without altering the system ability to perform gene expression.

## 1. Introduction

*In vitro* transcription translation systems have numerous applications, among them the study of gene expression and regulation, biochemical reactions and metabolic pathways (Dudley et al., 2015; Hold and Panke, 2009; Rolf et al., 2019; Schenkelberger et al., 2017). While *in vivo* the concentrations of different biochemical compounds can often only be estimated, *in vitro* systems allow for adjustable concentrations by addition or removal (Hold and Panke, 2009; Rolf et al., 2019; Sullivan et al., 2016). Optimal concentrations of their ingredients, e.g. for maximal protein yield or maximal reaction velocity, depend on the particular application (Noireaux et al., 2005; Shin and Noireaux, 2012).

The residue specific incorporation of non-canonical amino acids into proteins is a far reaching application of cell-free transcription-translation systems. Non-canonical amino acids as a surrogate at canonically prescribed locations can lead to new or improved properties of proteins with potential applications in biochemistry, biotechnology and biomedicine (Worst et al., 2016, 2015). However, for residue specific incorporation of non-canonical amino acids even small amounts of a corresponding canonical one will preferentially bind at the level of aminoacyl-tRNA-synthetases (Worst et al., 2016). Experiments by Hopfield have shown that a mechanism termed kinetic proofreading leads to a discrimination of up to 1/1000 against the incorrect amino acid (Hopfield, 1974; Hopfield et al., 1976).

*Escherichia coli* based expression systems are prepared “devoid” of amino acids, however, still there is a high level of remaining L-lysine, compared to other amino acids (Worst et al., 2016). In the following we will discuss methods for the removal of canonical amino acids, in particular L-lysine.

(Pure) expression systems contain only the strictly necessary compounds for gene expression. These compounds are individually expressed, purified and mixed at a later stage (Gao et al., 2019; Singh-Blom et al., 2014). This avoids the presence of residual matter from the cell lysate (in particular amino acids such as lysine). However, the usage of such systems is expensive (in time or money). Moreover results cannot be transferred to *in vivo* situations because of the minimalistic constitution of the pure system (Gao et al., 2019; Singh-Blom et al., 2014). The system depends on the bacteriophage T7 RNA polymerase, which leads to high protein yield. At the same time the system remains limited in terms of transcription regulation mechanisms (Shin and Noireaux, 2010).

Our cell-free expression systems are derived from *E. coli* cell lysates. They include the entire endogenous transcription-translation machinery (Garenne et al., 2019; Schenkelberger et al., 2017; Worst et al., 2015). In principle lysine-free expression systems could be derived from lysine auxotrophic *E. coli* strains (Gao et al., 2019; Singh-Blom et al., 2014). However, these strains require addition of the depleted amino acid for growth, leading again to the presence of residual amino acids in the final extract. Here, our aim is the creation of a method that will not affect extract preparation and that could be used for any cell-free system.

There are different methods to remove particular substances from an expression system. Filtering processes separate as a function of different properties, like hydrophobicity, size, charge, however, confounding lysine with other compounds with similar properties (Brödel et al., 2014; Gao et al., 2019). Antibodies or aptamers enable the separation of compounds in a much more specific way. To the best of our knowledge, however, there is no antibody against free lysine on the market. Related products will only recognize lysine in peptides and proteins (Steinbrecher et al., 1984; Xu et al., 2017). Furthermore antibodies are sensitive to chemical conditions (Jayasena, 1999; Song et al., 2008). A lysine aptamer can bind very specifically against lysine (Carter and Kataky, 2017; Fiegland et al., 2012; Sudarsan et al., 2003). However, depending on the type of aptamer its binding affinity is subject to change as a function of conditions. In our case the presence of K^+^ or Mg^2+^ may interfere (Carter and Kataky, 2017; Fiegland et al., 2012; Shin and Noireaux, 2012, 2010).

In this paper we describe a bead-based method for the removal of residual lysine from an all *Escherichia coli*-based cell-free expression system. Our method contains two steps. A first removal is obtained by the incorporation of lysine into a dummy protein with high lysine content. The second step uses the electrostatic attraction between lysine and DNA. The proposed method maintains the extract’s ability to perform gene expression.

## 2. Material and methods

Our cell-free expression system is prepared from *E. coli* as previously described (Shin and Noireaux, 2010; Sun et al., 2013).

The reporter plasmid is pBest-p15a-Or2-Or1-Pr-UTR1-deGFP-6xHis-T500. This plasmid is optimized for our cell-free system. It can be derived from the commercially available plasmid pBest-Or2-Or1-Pr-UTR1-deGFP-T500 (Addgene Cat# 40019) (Finkler and Ott, 2019; Schenkelberger et al., 2017; Shin and Noireaux, 2010). The gene for the high-lysine dummy protein (*HLDP*), DNA-sequence in Supplementary Fig. S1, is from Geneart (Thermo Scientific), subcloned into the pBest-p15a-Or2-Or1-Pr-UTR1-deGFP-T500 plasmid using the *BamH*I and *Xho*I restriction sites to obtain pBest-p15a-Or2-Or1-Pr-UTR1-deGFP-HLDP-T500. The deGFP sequence renders the expression of HLDP by production of the fluorescent fusion-protein deGFP-HLDP (GHLDP).

Adenine repeats impede gene expression (Arthur et al., 2015; Koutmou et al., 2015). Accordingly the sequence of the HLDP was chosen in such a way that lysine is at every second position, separated by any other canonical amino acid. The number of lysine in our HLDP was 53, 3 for all other amino acids (except histidine: six of them are used for His-tagging HLDP). The amino acid sequence can be written as M(XK)_53_H_6_, where X stands for all amino acids except lysine and histidine (3 of each amino acid, 2 methionines, see amino acid-sequence in Supplementary Fig. S1). While the production of this protein reduces the lysine content drastically, compared to lysine, the content of the other amino acids does not decrease significantly.

The gene coding for the fusion-protein GHLDP is linked to magnetic beads (Finkler and Ott, 2019; Invitrogen, 2017). Using the primers Biotin-TEG-Seq1 (TEG stands for TriEthyleneGlycol) (5′-Biotin-TEG-CAC CAT CAG CCA GAA AAC C-3′, Metabion) and Seq4 (5′-GAG CTG ACT GGG TTG AAG G-3′, Metabion) and streptavidin-coated magnetic beads T1 (Dynabeads® MyOne™ Streptavidin T1, Invitrogen) we obtain the construct T1-Biotin-TEG-Seq1-Or2-Or1-Pr-UTR1-deGFP-HLDP-T500-Seq4 (T1-*GHLDP*) at a DNA-concentration of 400 nM.

As previously shown, mRNA is quite stable in our system. It needs to be removed before new genes can be added (Finkler and Ott, 2019). For removal, DNA complementary to the mRNA of *GHLDP* is used. It also removes residual lysine. We use biotinylated single-stranded DNA complementary to the 3′-end of the Seq1-Or2-Or1-Pr-UTR1-deGFP-HLDP-T500-Seq4-mRNA (5′-Biotin-TEG-CGG CGG GCT TTG CTC GAG TTA GTG GTG ATG GTG ATG-3′, Metabion), linked to streptavidin-coated magnetic beads T1 (3’). Further we used poly-T-coated beads (poly-T) (Dynabeads™ Oligo(dT)25, Invitrogen), complementary to the poly(A) tail of the transcribed mRNA. Both ssDNA-bead stock solutions were at 5 mg/ml.

Experiments were performed on ice. The following ingredients were added to 30 μl of crude *E. coli* cell-free extract: 3.6 μl magnesium glutamate (at 100 mM), 2.7 μl potassium glutamate (at 3 M), 6.43 μl PGA-buffer (3-Phospho-Glyceric Acid) (14x), 4.5 μl 40% (v/v) PEG8000, 11.25 μl amino acids - lysine (mixture contains all canonical amino acids but no lysine at 6 mM except Leu at 5 mM), 3 μl GamS (Shortened lambda phage Gam protein, prevents linear DNA degradation (Garamella et al., 2016; Sun et al., 2014)) (at 99 μM) and 1.52 µl double-distilled water. The obtained 63 µl were aliquoted in 8.4 µl (seven samples in a batch).

For experiments 1.5 µl of the T1-*GHLDP* stock solution and 2.1 µl of double-distilled water were added to the 8.4 µl aliquots. This concentration is optimal for bead-bound DNA (Finkler and Ott, 2019). Aliquots filled up with double-distilled water to 12 µl were taken as a reference for comparison.

Samples were incubated at 29 °C for 30 minutes followed by magnetic separation of T1-*GHLDP* if present. During this time a mixture containing 50 μl of the 3′ and 50 μl of the poly-T stock solution was freshly prepared according to poly-T manufacturer’s protocol to obtain dried ssDNA coated beads (Ambion, 2012). After T1-*GHLDP* separation the supernatant was transferred to the prepared ssDNA coated beads and incubated for 15 minutes at room temperature at 230 rpm, followed by magnetic separation of the ssDNA coated beads. Subsequently 10 nM pBest-p15a-Or2-Or1-Pr-UTR1-deGFP-6xHis-T500 was added to the supernatant to visualize the incorporation of residual lysine in deGFP. To see if all these preparation steps affect the system’s ability to perform further gene expression, we also added 1 mM L-lysine to the supernatant with pBest-p15a-Or2-Or1-Pr-UTR1-deGFP-6xHis-T500 as reference. The increased error in this measurement (see Fig. 2) is due to small volume pipetting of highly concentrated lysine (about 0.3 µl of a 40 mM solution) required for maintaining overall concentration constant. For relative quantification of deGFP, 10 μl of samples were transferred to a microwell plate (Nunc™ 3 84-Well Optical Bottom Plates # 242764, Thermo Scientific) and the time course of fluorescence-intensity (see Fig. 2) was recorded with a microplate reader (POLARstar OPTIMA, BMG LABTECH) over 16 hours. Measurements with an initial *GHLDP* expression were corrected to obtain the fluorescence-intensity of the reporter protein deGFP only. The fluorescence of GHLDP was estimated by performing the corresponding measurement without addition of 10 nM pBest-p15a-Or2-Or1-Pr-UTR1-deGFP-6xHis-T500. The software Origin (Origin 2019b, OriginLab) was used for data evaluation.

It should be mentioned that using magnetic beads for our purpose requires very accurate processing. Using less beads results in diminished lysine removal, increased bead concentrations can remove other compounds in critical amounts so that system efficiency is reduced. Further, incomplete removal of magnetic beads from our expression reaction may interfere with gene expression. The amount of residual lysine differs between different stocks of crude extract. Therefore the amount of magnetic beads has to be adjusted for each stock. The amount of amino acids, magnesium glutamate and potassium glutamate must be adjusted for each stock for optimal cell-free protein synthesis (Finkler and Ott, 2019; Shin and Noireaux, 2010).

## 3. Results and discussion

As illustrated in Fig. 1 the first step towards lysine removal consists of its incorporation into a GHLDP (deGFP-high-lysine dummy protein, Fig. 1 A). For *GHLDP* expression all amino acids except lysine are added to the cell-free extract, resulting in lysine the limiting factor. GHLDP production can be observed by measuring its fluorescence-intensity. Because the expression time is short (30 minutes) not all of the residual lysine is incorporated into GHLDP. A prolonged *GHLDP* expression results in less residual lysine but also causes fatigue of the cell-free system. Fig. 1 B illustrates how we remove the remaining lysine through electrostatic attraction to DNA. Single-stranded DNA, complementary to the *GHLDP*-mRNA, is added to remove mRNA and stop *GHLDP* translation (and *GHLDP* expression) (Finkler and Ott, 2019; Naiser et al., 2008). Lysine is removed at the same time from the expression system. As a result the addition of DNA coding for deGFP does lead to significant production of deGFP only if lysine is added at the same time.

**Fig. 1.**
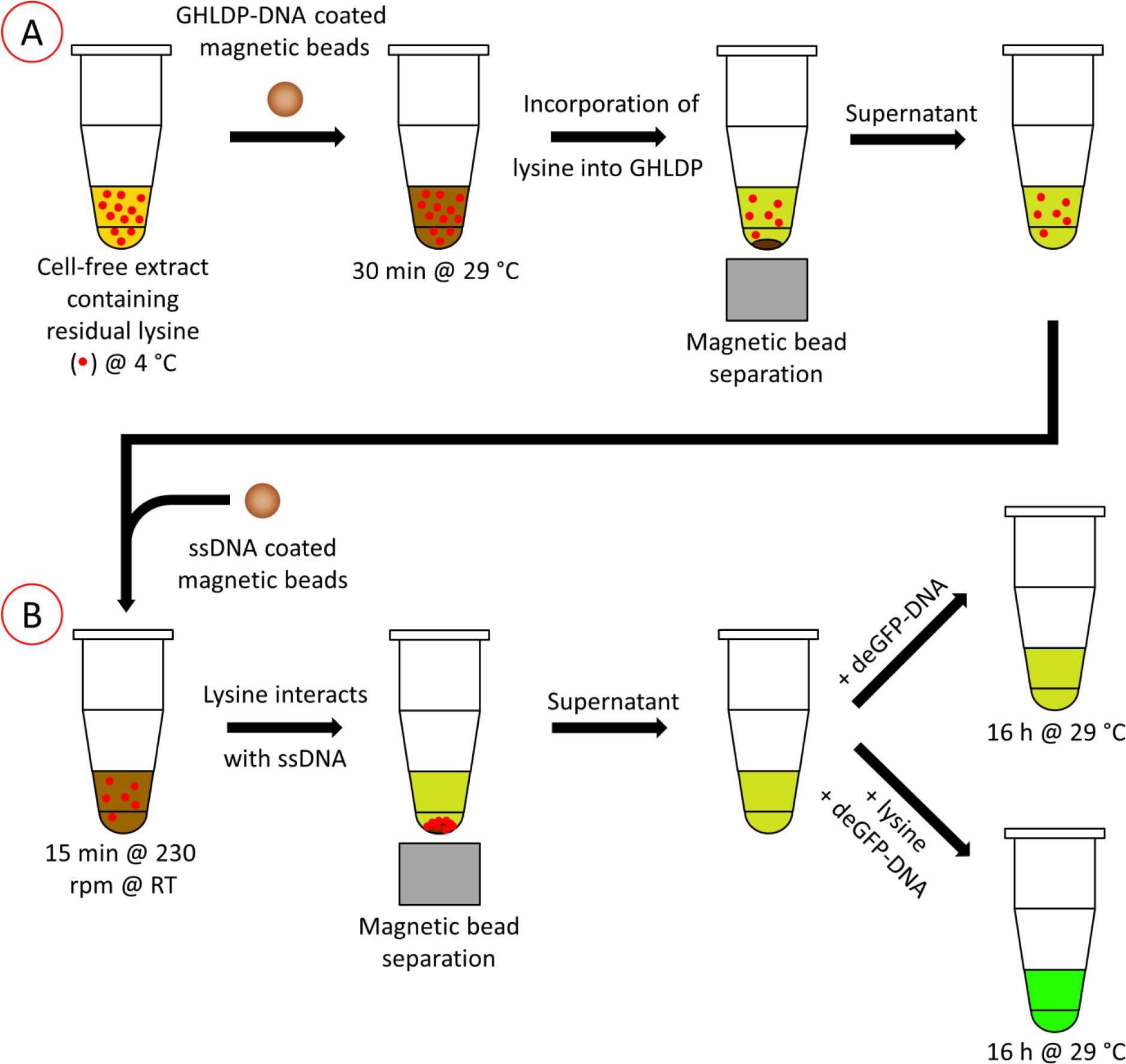
Scheme of the two step removal of residual lysine in a cell-free expression reaction. The process consists of A) lysine incorporation into a deGFP-high-lysine dummy protein (GHLDP) and B) removal of lysine through binding of lysine to DNA-coated beads. After the two step procedure, addition of deGFP-DNA will not lead to expression because of missing lysine. Only if lysine is added, deGFP will be produced.

Fig. 2 shows the measured time course of fluorescence-intensity of expressed deGFP over 16 hours for different samples.

**Fig. 2.**
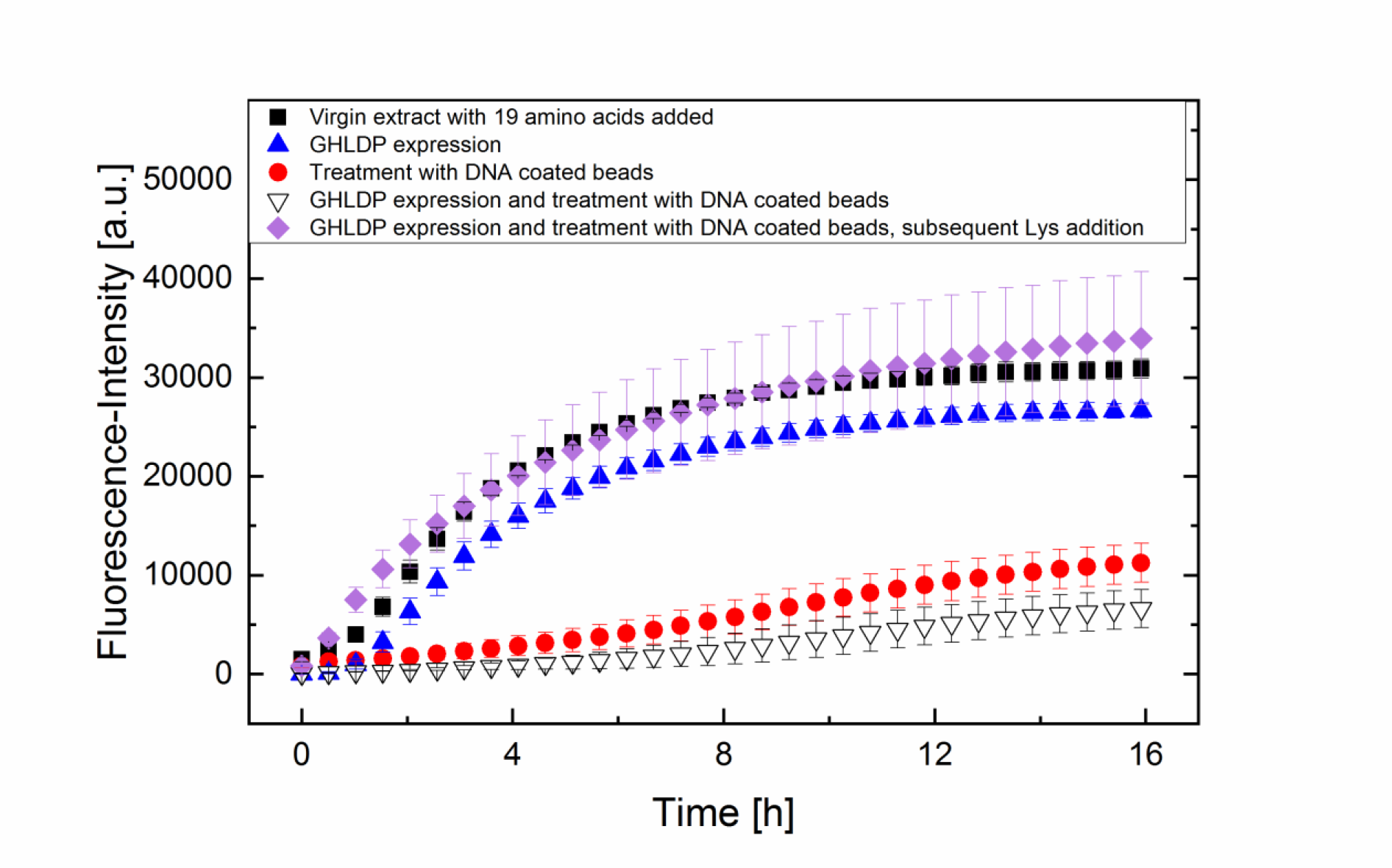
Time course of deGFP fluorescence-intensity from different cell-free expression reactions. Initial addition of 20 amino acids to a virgin extract results in a final deGFP fluorescence-intensity of about 75000 ± 5000 a.u. (data not shown). All reactions shown were performed with initial addition of only 19 amino acids (all except lysine), making the residual lysine from the initial preparation the limiting factor. Black squares represent the time course of deGFP fluorescence-intensity from a virgin 19 amino acid extract. The other studied samples differ by their treatment to remove residual lysine. The incorporation of residual lysine into GHLDP (blue triangles, following Fig. 1 A), or lysine separation by treatment with ssDNA coated beads (red circles, following Fig. 1 B), results in lower lysine content. The combination of both methods exhibits a cumulative effect (black open triangles). The subsequent addition of lysine mediates the production of deGFP (purple diamonds) comparable to the virgin extract with 19 amino acids added.

To check for self-similarity of the curves for lysine deprived systems, we rescaled the time and fluorescence-intensity (characteristic time, τ, and characteristic amplitude, A). The scaling values are listed in table 1.

**Table 1.** Characteristic time until inflection of the time-resolved fluorescence-intensity (Characteristic time, τ) and characteristic amplitude of fluorescence-intensity (Characteristic amplitude, A) for different samples. These values were used for rescaling the curves in Fig. 2 (Fig. 3).

| Sample | Characteristic time, $\tau$ [a.u.] | Characteristic amplitude, $A$ [a.u.] |
| --- | --- | --- |
| Virgin extract with 19 amino acids added | 1 | 1 |
| *GHLDP* expression | 1.1 | 0.91 |
| Treatment with DNA coated beads | 3.29 | 0.5 |
| *GHLDP* expression and treatment with DNA coated beads | 4.06 | 0.37 |

By rescaling the sigmoid curves with the characteristic parameters in table 1 we obtained Fig. 3.

**Fig. 3.**
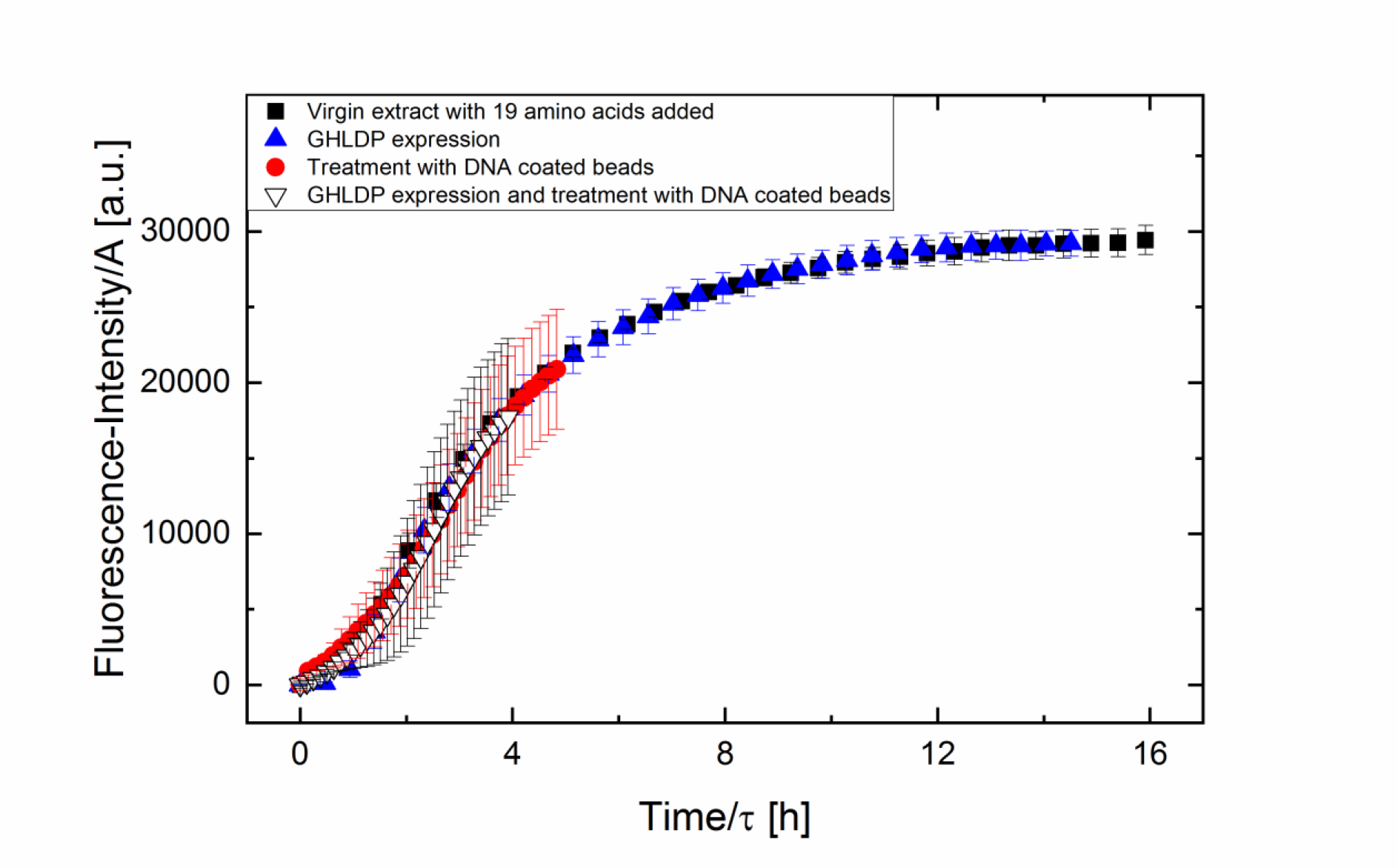
Rescaled curves from Fig. 2, using the parameters in table 1.

According to Fig. 3, the curves are self-similar, suggesting a power law description A ̴ τ^γ^. Taking γ as a fitting parameter, we obtain γ = −0,64 ± 0,06. A value of −1 corresponds to a simple diffusion-limited process. The result shows that other processes come into play. The exponent is larger than −1, compatible with the idea that proteolysis releases lysine, yielding increased amounts of deGFP. To investigate this idea, we performed experiments with protease inhibitors (Protease Inhibitor Cocktail P8849, Sigma-Aldrich). High concentrations resulted in a non-functioning expression system (possibly non-specific side effects), while lower concentrations had no effect. We conclude that proteolysis is unlikely. In experiments we observed that our expression reaction can be kept for at least 1 hour without fatigue if no expression is mediated by DNA addition. Hence another possibility for the observed scaling exponent could be due to system fatigue caused by expression. This would lead to more efficient usage of lysine at low expression rates. The asymptotic endpoints of deGFP expression (the amplitude) then represent a lower bond for lysine removal. Accordingly we interpret A⋅[lysine]_Virgin extract_ as an upper bound of the remaining lysine and τ^−1^⋅[lysine]_Virgin extract_ as an estimation of the lysine concentration.

Because lysine is the limiting factor in these experiments, we can interpret the deGFP fluorescence-intensity after 16 hours of expression to reflect the initial content of lysine in the system (Shin and Noireaux, 2010). With this assumption, the incorporation of residual lysine into GHLDP decreases the lysine content by 14 % while lysine removal through DNA coated beads resulted in a 64 % reduction. The combination of both processes reduces the initial lysine content by almost 80 %. This is in excellent agreement with the values for τ.

According to literature, with our system and all amino acids provided, under optimal conditions a deGFP concentration of 0.65 mg/ml is reached (Shin and Noireaux, 2010). We deduce a concentration of 0.025 mg/ml of residual lysine in our virgin extract. Accordingly the 12 µl of our expression reaction contain at least about 300 ng of residual lysine. Our two step method decreased this to about 60 ng.

Fig. 2 shows that the difference in the final fluorescence-intensity between the black squares and the blue triangles (4636,24 ± 1925,96 a.u.) is similar to the difference between the red circles and the black open triangles (4250,33 ± 914,91 a.u.). In both cases the difference in the final fluorescence-intensity was obtained by initial *GHLDP* expression. Although additional treatment with DNA coated beads decreases the final fluorescence-intensity, the relative contribution of GHLDP to the decrease is the same as without the additional bead treatment. The two steps of our proposed method to remove residual lysine are cumulative.

Subsequent addition of lysine after initial two-step removal of residual lysine led to strong deGFP expression (Fig. 2, purple diamonds). This shows that our proposed method is minimally invasive. The expression system is not affected in its ability to perform gene expression. The subsequent addition of lysine to an expression reaction whose lysine was removed with our two step method (Fig. 2, purple diamonds) led to only the half the amount of deGFP, compared to adding 20 amino acids to a virgin extract. Beginning fatigue of the expression system by initial *GHLDP* expression is one possible explanation.

We achieved a five-fold terminal fluorescence-intensity by addition of lysine at t = 0 h to an extract that was initially treated with our two step method (Fig. 2, black open triangles and purple diamonds), which means that 80 % of the produced deGFP is from the added (not the residual) lysine.

## 4. Conclusion

We presented a new method, simple and fast, to remove a huge fraction (80 %) of residual lysine from an *E. coli* based cell-free expression system. Our minimally-invasive method does not alter the system in its ability to perform gene expression. After lysine removal the subsequent addition of deGFP-DNA and lysine resulted in strong deGFP production.

We found a relationship between the characteristic amplitude, A and the characteristic time, τ in experiments with limited lysine content, A ̴ τ^−0.64±0.06^. Because the exponent differs from −1 we suggest that other processes, like expression mediated system fatigue may play a role.

Although our method removes up to 80 % of the residual lysine, different experiments may require to even further decrease the amount of lysine. To achieve this, modifications in the crude extract preparation like prolonged dialysis (from 3 h (Shin and Noireaux, 2010) to up to 12 h) may be useful. Further the usage of a lysine auxotrophic bacterial strain for extract preparation in combination with our method may be an effective way to decrease the residual lysine to even lower levels.

The indirect removal of lysine by incorporation into a dummy-protein can be transferred to other amino acids easily. In contrast, the removal of amino acids through binding to DNA is unlikely to work for negatively charged amino acids.

## Author contributions

AO and MF designed the research. MF performed the research with the help of ÖK and FG. MF analyzed data with the help of AO, ÖK and PH. MF and AO wrote the paper.

## Acknowledgements

This work was supported by the collaborative research center SFB 1027 funded by Deutsche Forschungsgemeinschaft (DFG) and by the Human Frontier Science Program (HFSP, RGP0037/2015). We thank Emanuel Worst for help and fruitful discussions.

