## Supplementary material for "A bead-based method for the removal of the amino acid lysine from cell-free transcription-translation systems"

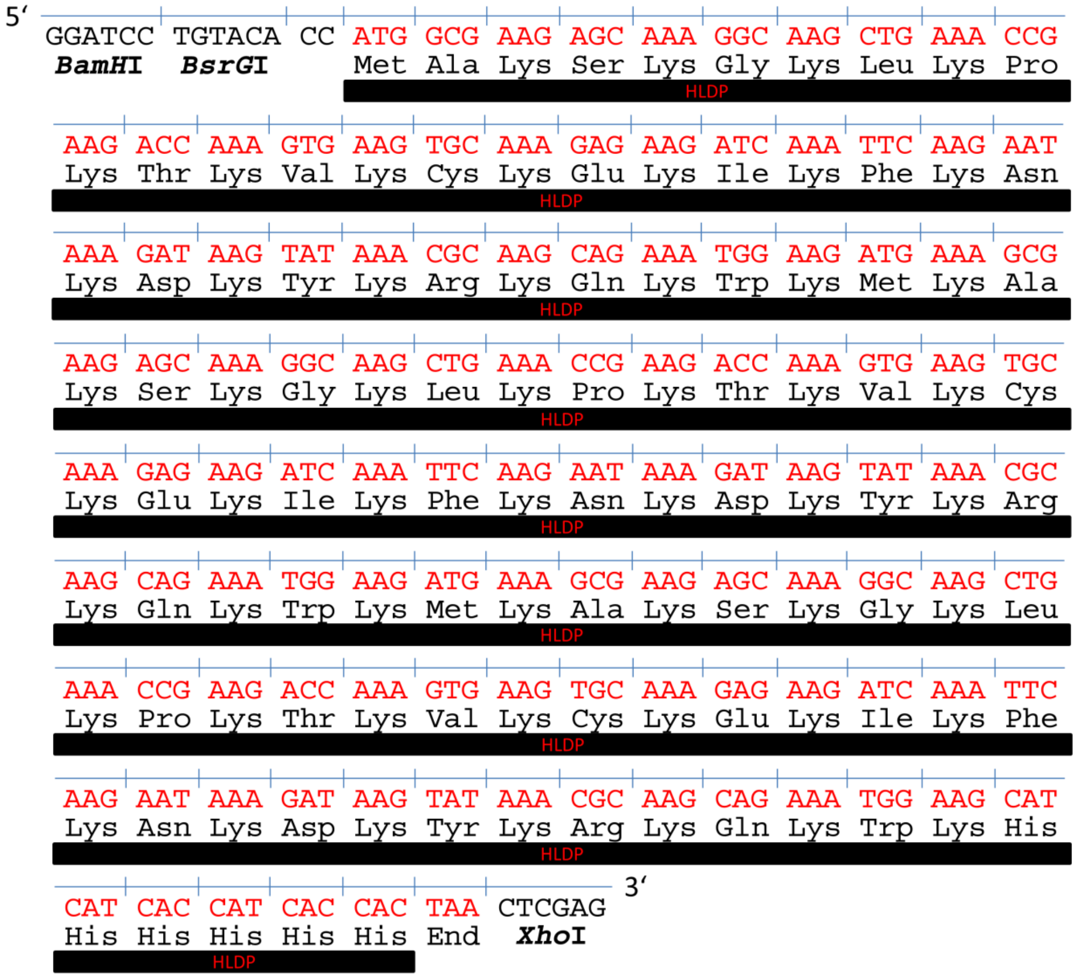


**Fig. S1. DNA- and the deduced amino acid-sequence of HLDP. Because the gene expression of a polylysine gene was not successful if adenine repeats were present (Arthur et al., 2015; Koutmou et al., 2015) the sequence was adapted to a maximum of 3 adenines in a row. The codons coding for lysine are AAA and AAG, and the insertion of another amino acid between two lysines was necessary to prevent adenine stretches. Codons optimal for our system were chosen using the Kazusa codon usage database preventing 4 adenines in a row. Alanine was chosen to be a spacer between the initial methionine and the first lysine. Further a 6xHis-Tag was added to the last lysine. To keep the content of all amino acids except lysine and histidine constant, each amino acid is present at the same number of 3. This gives 53 lysine residues in our HLDP. *BamH*I and *Xho*I restriction sites are flanking our sequence.**
